# Proinflammatory stress activates neutral sphingomyelinase 2 based generation of a ceramide-enriched β cell EV subpopulation

**DOI:** 10.1101/2024.04.17.589943

**Authors:** Jerry Xu, Arianna Harris-Kawano, Jacob R. Enriquez, Raghavendra G. Mirmira, Emily K. Sims

## Abstract

β cell extracellular vesicles (EVs) play a role as paracrine effectors in islet health, yet mechanisms connecting β cell stress to changes in EV cargo and potential impacts on diabetes remain poorly defined. We hypothesized that β cell inflammatory stress engages neutral sphingomyelinase 2 (nSMase2)-dependent EV formation pathways, generating ceramide-enriched EVs that could impact surrounding β cells. Consistent with this, proinflammatory cytokine treatment of INS-1 β cells and human islets concurrently increased β cell nSMase2 and ceramide expression, as well as EV ceramide staining. Direct chemical activation or genetic knockdown of nSMase2, or treatment with a GLP-1 receptor agonist also modulated cellular and EV ceramide. Small RNA sequencing of ceramide-enriched EVs identified a distinct set of miRNAs linked to β cell function and identity. Coculture experiments using CD9-GFP tagged INS-1 cell EVs demonstrated that either cytokine treatment or chemical nSMase2 activation increased EV transfer to recipient cells. Children with recent-onset T1D showed no abnormalities in circulating ceramide-enriched EVs, suggesting a localized, rather than systemic phenomenon. These findings highlight nSMase2 as a regulator of β cell EV cargo and identify ceramide-enriched EV populations as a contributor to EV-related paracrine signaling under conditions of β cell inflammatory stress.

**Article Highlights:** ***a.** Why did we undertake this study?:* Mechanisms connecting β cell stress to changes in extracellular vesicle (EV) cargo and potential impacts on diabetes are poorly defined.

***b.** What is the specific question we wanted to answer?:* Does β cell inflammatory stress engage neutral sphingomyelinase 2 (nSMase2)-dependent EV formation pathways to generate ceramide-enriched EVs.

***c.** What did we find?:* Proinflammatory cytokine treatment of β cells increased β cell ceramide expression, along with EV ceramide in part via increases in nSMase2. Ceramide-enriched EVs housed a distinct set of miRNAs linked to insulin signaling. Both cytokine treatment and nSMase2 activation increase EV transfer to other β cells.

***d.** What are the implications of our findings?:* Our findings highlight nSMase2 as a regulator of β cell EV cargo and identify ceramide-enriched EV populations as a contributor to EV-related paracrine signaling under conditions of β cell inflammatory stress.

## Introduction

Pancreatic β cell failure is the cornerstone of diabetes development, and β cells may not be innocent bystanders in this process (1). Rather, β cells may exhibit maladaptive responses to inflammatory stress in different forms of diabetes, such as prolonged exposure to systemic elevations in proinflammatory cytokines and saturated free fatty acids in type 2 diabetes (T2D) or autoimmune attack of β cells and cytokine exposure in type 1 diabetes (T1D) (2, 3). Specifically, metabolic stress pathways intrinsic to the β cell can hasten the progression to apoptosis or enhance β cell responsiveness to inflammatory or immune-mediated destruction (2, 4). Clear delineation of the contributions of intrinsic β cell dysfunction to diabetes development is critical, as it may ultimately pave the way for novel β cell-targeted therapies and novel biomarkers of disease.

Extracellular vesicles (EVs) are membrane bound nanoparticles housing molecular cargo (5). EVs can be transferred to or interact with other cells as a means of cell:cell communication (6–8). The most abundant β cell EVs are small EVs, predominantly comprised of exosomes, or EVs formed from exocytosis of multivesicular endosomes (MVEs) (5, 8). β cell EV cargo change under stress and disease conditions, and some data suggest β cell EVs may serve as paracrine effectors in the islet microenvironment in health and disease (6, 7, 9–16). However, mechanisms of changes in β cell EV cargo remain poorly defined.

In non-islet cells, diverse molecular mechanisms have been identified as regulating EV biogenesis and cargo loading, including ceramide-dependent MVE vesicle formation and release of ceramide-enriched small EV subpopulations (17–20). Ceramides are bioactive sphingolipids, molecules containing a sphingoid base backbone attached to a fatty acid side chain that can function as membrane signaling components in stress responses (21). Ceramide generation is linked to several β cell stress pathways, including oxidative stress, ER stress, and pro-apoptotic signaling (22–28). Based on this linkage, we hypothesized that islet inflammatory stress and increased ceramide generation would increase in ceramide-enriched EV populations. We specifically sought to test whether this occurs due to increases in neutral sphingomyelinase 2 (nSMase2), an enzyme that hydrolyzes sphingomyelin to generate ceramide and has been linked to changes in EV microRNA (miRNA) cargo in other systems (17, 18, 29).

## Research Design and Methods

### Culture of Cells and Human Islets

Specific reagents utilized are detailed in **Supplemental Table 1**. β cell lines, including INS-1 823/13 and MIN6 (originally obtained from Chris Newgard, and J. Miyazaki), were cultured as described previously (8, 30), but with 10% exosome depleted FBS (purchased or prepared by ultracentrifugation at 120,000 xg for 20 h at 4°C followed by filtration (0.22 µm PES filter). Human islets and dissociated islet cells were cultured in human islet culture medium DMEM plus 10% FBS. Human islets were provided from the Integrated Islet Distribution Program at City of Hope and The Alberta Diabetes Institute IsletCore (**Supplemental tables 2 and 3**).

To model the proinflammatory milieu of developing diabetes, cells were exposed to IL1β (5 - 10 ng/mL) or a cytokine mix of IL1β (5 ng/mL), TNFα (10 ng/mL), and IFNγ (100 ng/mL) for 24-48 hours as described (8, 30). To model β cell endoplasmic reticulum stress and DNA damage, cells were exposed to 24 hours of 5 nM thapsigargin, 1 µM tunicamycin or or 50 nM doxorubicin. To chemically induce nSMase 2 activation, cells were treated with 24 hours 5 µM Caffeic acid phenethyl ester (CAPE). To examine the effect of exendin-4, we pre-treated INS-1 cells with 10LJ nM exendin-4 or vehicle for 15 min, then treated cells with IL1β in the presence of 10LJ nM exendin-4 or vehicle for another 24 h.

### Generation of INS-1 Derivative Cell Lines

INS-1 cells with green fluorescent protein (GFP)-tagged CD9, an exosome surface marker, were generated by amplification of CD9, followed by amplicon cloning into a pLJM1-EGFP lentiviral vector (Addgene plasmid #19319) to express CD9 as a C-terminal-tagged GFP fusion protein, using circular polymerase extension cloning. Stable INS-1 cell lines expressing mCherry fluorescent protein, INS-1 nSMase2 knockdown cells, INS-1 scramble shRNA cells were generated by transfecting plasmids with mCherry in pReceiver-Lv213 vector, rat nSMase2 shRNA with GFP reporter, and a non-effective 29-mer scrambled shRNA cassette in a pGFP-C-shLenti Vector, respectively. Transfected cells were selected with puromycin, pooled and sorted using fluorescence-activated cell sorting (FACS, BD FACSAria) into 96-well plates by the Indiana University Flow Cytometry Core. Individual cell colonies were isolated and expanded. mCherry expression and nSMase2 knockdown level were validated using flow cytometry and immunoblotting.

### EV Isolation

EVs were isolation using ultracentrifugation, affinity purification with biotinylated tetraspanin antibodies, or size exclusion chromatography (SEC). For ultracentrifugation, conditioned media was centrifuged at 3000g for 10 min to remove debris, filtered through 0.22 µm PVDF or PES filter to remove large vesicles, then centrifuged at 120,000g for 1.5 h at 4°C. Pelleted EVs were washed and centrifuged in 120,000g for 1.5 h at 4°C then suspended in 100 µl of 0.22 µm filtered PBS.

For immunoaffinity isolation, filtered media was concentrated using a 100K MWCO concentrator. EVs were captured by magnetic streptavidin beads coupled with biotinylated tetraspanin antibodies (CD9, CD63, CD81) using Exo-Flow Capture kits according to the manufacturer’s manual. For use with capture kits, we biotinylated 100 µg mouse IgM isotype control or mouse monoclonal anti-ceramide antibody with an antibody biotinylation kit for immunoprecipitation following manufacturer’s instructions. To selectively capture ceramide EVs, we coupled streptavidin magnetic beads with biotinylated ceramide antibody as above. Captured EVs were stained with Exo-FITC exosome stain and quantified using flow cytometry.

Human plasma EVs were purified by SEC using qEVsingle /35 nm Legacy columns following the manufacturer’s instructions.

For EV validation, nanoparticle tracking analysis (NTA) was performed using ZetaView (ParticleMetrix GmbH, Ammersee, Germany) with 100 nm microspheres as particle size standards. For Transmission Electron microscopy (TEM), isolated EVs were eluted with exosome elution buffer, washed with PBS, and desalted and concentrated, then fixed with electron microscopic fixation buffer, sent to and imaged by the Electronic Microscopy Core at University of Nebraska Medical Center.

### Coculture Experiments

CD9GFP EV fluorescence was validated using flow cytometry (**Supplemental figure 1A**). Recipient INS1-mCherry cells and donor INS1-CD9GFP cells were mixed at the ratio of 1:1 and seeded in 6-well plate or dishes in INS1 culture medium. After incubating for 4 h, culture medium was replaced with exosome depleted FBS medium. Another 48 h or 72 h, the cells were dissociated by trypsinization and subjected to flow cytometry analysis to quantify the fluorescent intensity of recipient INS1-mCherry cells that internalized EVs carrying CD9GFP. EV transfer was also quantified after seeding INS1-CD9GFP cells onto 6-well plates, with wild-type INS1 cells in trans-well inserts for 48 or 72 hrs.

### Western Blotting

Immunoblots were performed as described (30). Blots were probed against anti-nSMase2 antibody (1:200), and anti-β-actin (1:1000). Goat anti-rabbit IgG or goat anti-mouse IgG were used as secondary antibodies and visualized using an Odyssey Fc imaging system (Li-Cor Biosciences).

### Flow Cytometry

INS-1 or MIN6 cells were harvested and filtered through 40 µm cell strainers to obtain single cell suspension. To analyze cell surface ceramides, cells were blocked and stained in blocking buffer 5% FBS/0.1% BSA/PBS/2 mM EDTA. To examine total cellular ceramides, cells were fixed in 4% methanol-free formaldehyde solution and permeabilized in 0.1% saponin/PBS/2 mM EDTA. Cells were blocked in the above blocking buffer plus 0.1% saponin. Cells were stained with monoclonal anti-ceramide antibody (1 µg/mL) and/or N-SMase2 antibody (1 µg/mL), and corresponding fluorescent conjugate second antibody (1 µg/mL). The monoclonal anti-ceramide antibody (31) (clone MID 15B4, MilliporeSigma) used recognizes free and bound ceramides and does not cross-react with sphingomyelin and other phospholipids in vitro and in vivo physiological conditions. Cells were resuspended in 2% FBS/PBS/2mM EDTA for flow cytometry analysis. Intact human islets were dispersed using 0.25% trypsin and neutralized with 10% in PBS, or by using enzyme-free cell dissociation buffer, filtered through a 40 µm cell strainer. Cells were fixed with 4% methanol-free paraformaldehyde and stained with anti-ceramide antibody (1 µg/mL) and/or N-SMase2 antibody (1 µg/mL) as above. Flow cytometry data were interpreted using FlowJo (Becton, Dickinson & Company).

For EV analysis, EV populations were captured using immunoaffinity and stained with anti-ceramide antibody (1 µg/mL) or isotype control for 1 h at 4°C, and washed x 3 with bead wash buffer, then further stained with fluorescent conjugated secondary antibody (1 µg/mL) for 30 min at 4°C, washed x3, then resuspended in 500 µL bead wash buffer for analysis.

### ELISA of Human Plasma EVs

Deidentified plasma from the IU CDMD biobank was utilized for testing of samples from children with type 1 diabetes (clinically diagnosed by a pediatric endocrinologist) or euglycemic controls matched for age, sex, and body mass index). Ethical approval was obtained from the IU Institutional Review Board and all participants provided informed consent or assent as appropriate and consistent with principles expressed in the Declaration of Helsinki. Collected EVs by SEC were lysed using 5x ELISA lysis buffer and diluted 1:80. Ceramide concentration was quantified using ELISA with Ceramide ELISA Kit or flow cytometry as above.

### RNA Sequencing and Expression Analysis

#### Single Cell RNA-Sequencing Analysis

To examine *Smpd3* expression, we used the mouse islet single cell atlas previously compiled (32). We downloaded the atlas from the cellxgene instance (https://cellxgene.cziscience.com/collections/296237e2-393d-4e31-b590-b03f74ac5070) and inputed the .rds format into Seurat (33) (v4.3.0) with R (v4.2.2). We then subsetted the metadata by disease state selecting only “normal”, “type 1 diabetes mellitus”, and “type 2 diabetes mellitus”; then further subsetted the atlas to only “beta” cells. Visualization was performed using VlnPlot functions and differential expression was performed with the FindAllMarkers function.

#### EV Small RNA Sequencing

Affinity isolated EVs were resuspended in PBS. Total RNA was isolated using miRNeasy Isolation Kits and on-column digestion of DNA with RNase-Free DNase Set. miRNA sequencing (RNA-seq) and analysis was performed by IU Center for Medical Genomics using platform NextSeq500. Differential miRNA expression analysis was performed with edgeR (34), using negative binomial generalized linear models with likelihood ratio tests. The leading fold changes were calculated as the average (root-mean-square) of the largest absolute log-fold changes between each pair of samples. Network analysis was performed using QIAGEN Ingenuity Pathway Analysis (IPA).

### Statistics

Data are plotted as mean +/-standard error. Statistical significance was determined using student’s two tailed t-test in GraphPad Prism. A p value of <=0.05 was considered statistically significant.

### Data and Resource Availability Statement

Data sharing and resource sharing are available upon reasonable request to authors.

## RESULTS

### β cell nSMase2 and ceramide, as well as β cell EV ceramide content, are increased under conditions of proinflammatory cytokine exposure

To examine the expression of sphingomyelin phosphodiesterase 3 (*Smpd3)*, which encodes nSMase2, we first examined published data on β cell RNA expression from the mouse islet single-cell atlas (32). Grouping by disease state, shown in **Figure 1A**, identified a significant (p<0.01) upregulation of *Smpd3* in β-cells from models of type 1 diabetes. A smaller increase appeared to be present in β cells from a model of type 2 diabetes, but this was not statistically significant. Based on these findings, we tested if proinflammatory cytokine exposure directly induces increases in nSMase2 by quantifying changes in protein levels. Using immunoblot and flow cytometry (**Figure 1B-C**), we observed that 24-hour IL1β treatment of INS-1 cells increased nSMase2 expression. In concert with changes in nSMase2, IL1β treatment significantly increased both total cellular ceramide expression (**Figure 1D**) and cell surface ceramide expression (**Figure 1E**). Similar patterns were also present in MIN6 cells after 24 h treatment with proinflammatory cytokine mix (**Supplemental Figure 1**). Findings were also verified in human islets treated with cytokine mix (**Figure 1F and 1G**).

**Figure 1.**
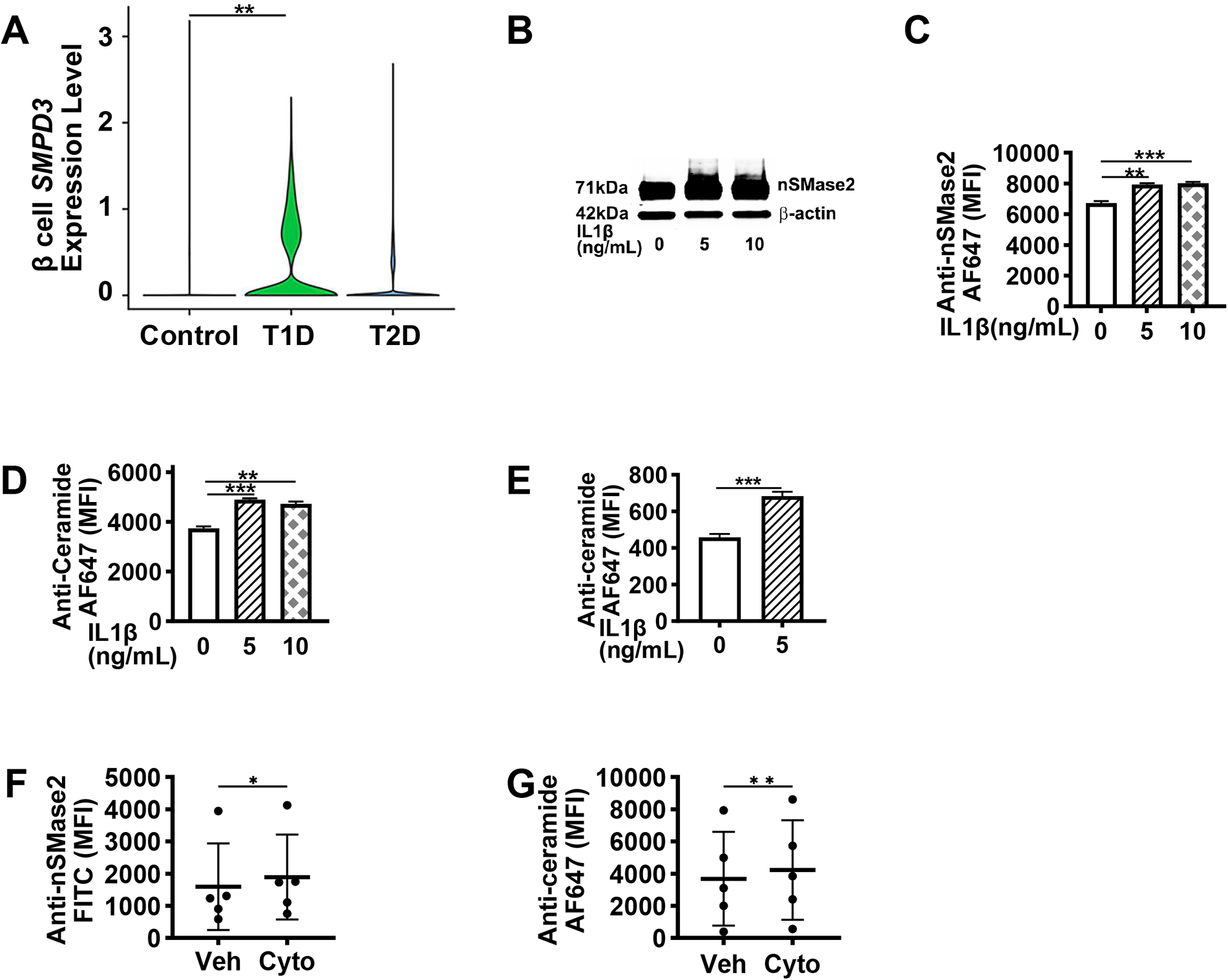
β cell nSMase2 expression and ceramide production are increased in concert under conditions of type 1 diabetes (T1D) and cytokine exposure. (A) Violin plots of β-cell *Smpd3* expression in control, T1D, and T2D models show a significant upregulation in β-cells from a mouse model of T1D (p<0.01). (B-E): In INS1 β cells, 24 hr IL1β (5 or 10 ng/mL) increases nSMase2 as measured via (B) immunoblot or (C) flow cytometry staining, as well as (D) total cellular ceramides and (E) surface ceramide flow cytometry staining. (F--G): Human islets were treated with 48 h mix of IL1β (5 ng/mL), TNFα (10 ng/mL) and IFNγ (100ng/mL), then dispersed for flow cytometry-based quantification of (E) ceramides and (F) nSMase2 in triplicate, for each of the indicated donors. INS1 n=9, with 3 separate experiments performed; human islets tested in triplicate, using samples from 5 individual donors. Data expressed as mean ± SEM; *p<0.05; **p<0.01; ***p<0.001

To understand if cytokine-induced increases in β cell nSMase 2 and ceramide were also associated with increases in EV ceramide, we performed tetraspanin immunoaffinity based small EV pull downs and quantified INS-1 and human islet small EV ceramides using flow cytometry. Small EV isolations were validated using TEM and NTA (**Figure 2A and 2B**) (35). For both INS-1 cells and human islets, treatment with IL1β or cytokine mix also increased small EV ceramide content (**Figure 2C and 2D**).

**Figure 2.**
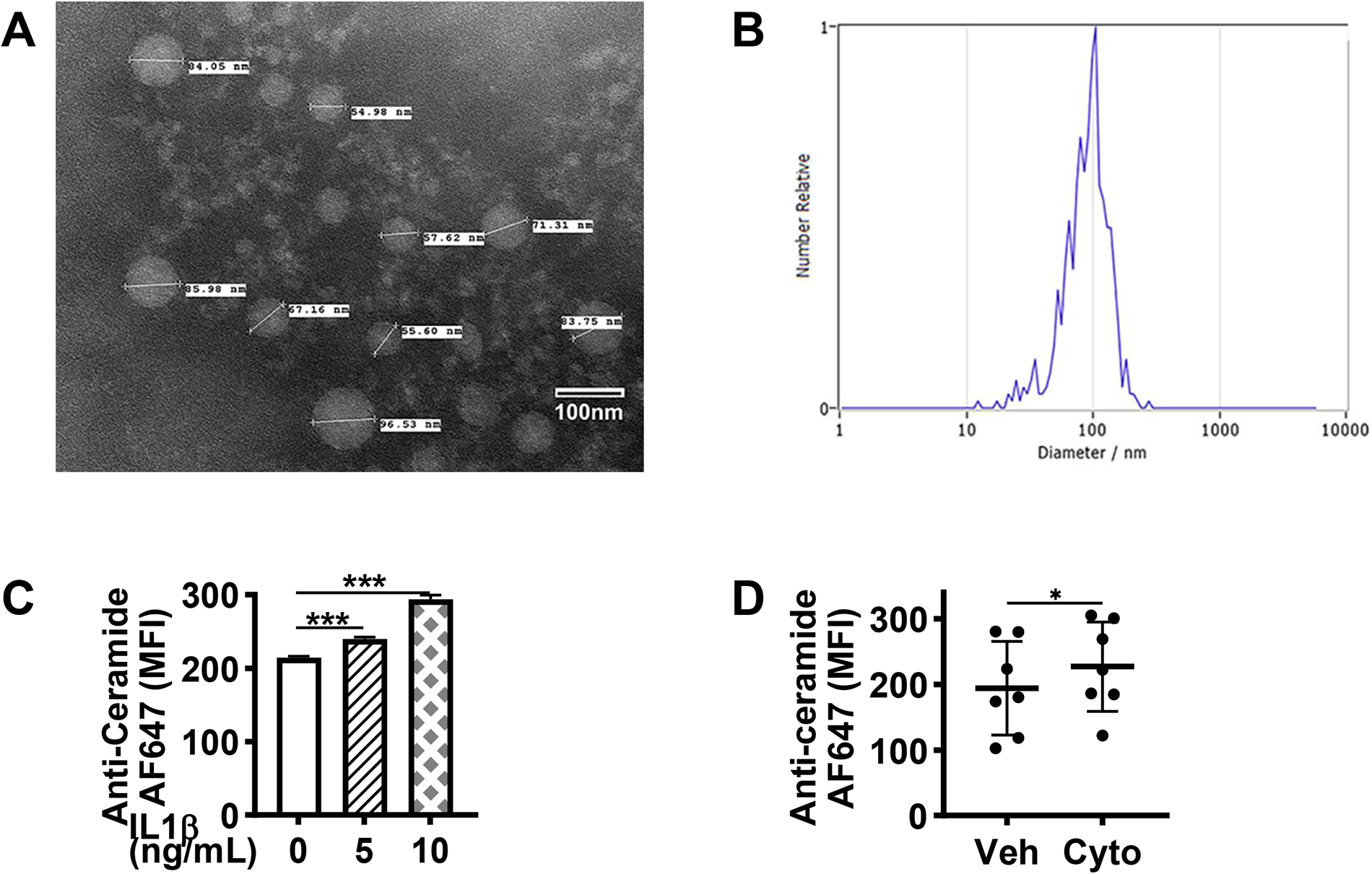
In concert with cellular nSMase2 and ceramide, cytokine treatment increases β cell EV ceramides. β cell small EVs were isolated using bead based tetraspanin antibody pulldown, eluted, washed, concentrated and verified using (A) electronic microscopy (direct mag. 110000x) and (B) nanoparticle tracking analysis. Flow cytometry-based staining for ceramide was performed to assess EV ceramide content in control and cytokine treated (C) INS1 cells and (D) human islets. INS1 n=9, with 3 separate experiments performed; human islets, triplicate, with 7 separate experiments performed. Data expressed as mean ± SEM; *p<0.05; **p<0.01; ***p<0.001

### Direct manipulation of cellular nSMase 2 impact EV ceramide content

Next, we aimed to directly test the impact of cytokine-induced increases in nSMase 2 expression on ceramides in cellular and small EV ceramide content. First, we exposed INS-1 cells to 24 h treatment with caffeic acid phenethyl ester (CAPE), a chemical inducer of nSMase2 activity and intracellular ceramide accumulation (36). CAPE treatment increased cell ceramides both at baseline and in response to IL1β treatment (**Figure 3A**). In parallel, treatment increased the total small EV number (**Figure 3B**) as well as small EV ceramide content (**Figure 3C**), suggesting that nSMase2 based increases in small EV ceramides may not only yield increases in individual EV ceramide cargo but also lead to increases in ceramide-enriched small EV populations.

**Figure 3.**
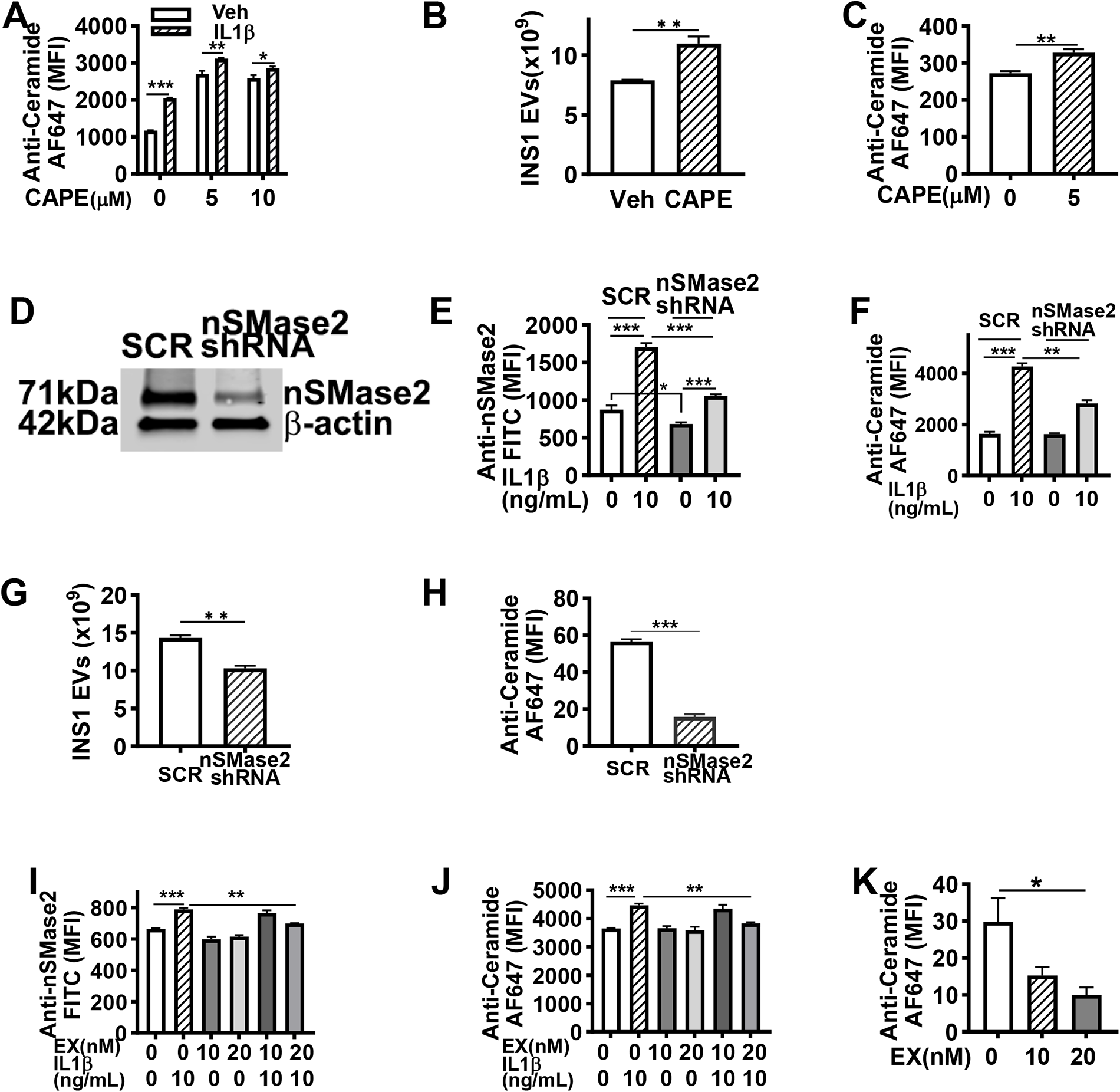
Modulation of nSMase2 activity impacts β cell ceramide-enriched EV generation. A-C: INS1 β cells were treated with 24 h of Caffeic acid phenethyl ester (CAPE) (5 µM), which increased cell ceramides (A), total EV secretion (B), as well as EV ceramide staining (C) with or without concurrent cytokine treatment. D) stable nSMase2 knockdown INS1 β cells were developed with rat nSMase2shRNA to reduce cellular nSMase2 levels at baseline and (E) in response to cytokine (10 ng/mL IL1β). (F) nSMase2 knockdown also reduced cytokine induced increases in cellular ceramide staining. (G, H) nSMase2 knockdown reduced total EV secretion and drastically reduced ceramide enriched EVs. (I-K): INS1 β cells were treated with the diabetes therapeutic and GLP-1 (glucagon-like peptide-1 receptor) agonist Exendin-4 at 20 nM for 24 h. Exendin-4 treatment (20 nM) significantly abrogated IL1β-induced cellular (I) nSMase2 and (J) at 20 nM, cellular ceramide staining. (K) Exendin-4 also significantly decreased EV ceramide staining. n=9, with 3 separate experiments performed.Data expressed as mean ± SEM; *p<0.05; **p<0.01; ***p<0.001

To assess the impact of reduced nSMase2 activity, we performed genetic knockdown (KD) of nSMase2 in INS-1 cells using shRNA (**Figure 3D**). nSMase2 KD cells exhibited reduced expression of nSMase2 both at baseline and in response to cytokine treatment (**Figure 3E**). Consistent with a direct effect of nSMase2 activity on cytokine-induced increases in cellular ceramides, cellular ceramides were not reduced at baseline, but cytokine-induced increases in cellular ceramides were abrogated in nSMase2 KD cells (**Figure 3F**). In contrast to CAPE induced nSMase2 activation, nSMase2 KD yielded a reduced total small EV number (**Figure 3G**). This was associated with dramatic reductions in ceramide-staining in small EV ceramides on flow cytometry (**Figure 3H**), again supporting the idea that nSMase2 is linked to generation of ceramide-enriched small EV populations.

### Modulation of other intrinsic islet signaling pathways also impacts β cell EV ceramide content

Next, we applied treatment with exendin-4, a glucagon-like peptide 1 (GLP-1) agonist, which in addition to positive impacts on beta cell health and potentiation of insulin secretion has been shown to inhibit activation of de novo ceramide synthesis (37, 38). As shown in Figure **3I and 3J**, exendin 4 treatment did not impact INS-1 cellular nSMase2 and ceramide content at baseline but reduced IL1β -induced increases in cellular nSMase2 and ceramide content. Additionally, exendin-4 treatment yielded a dose-dependent reduction in EV ceramide content (**Figure 3K**).

To test the impacts of activation of other β cell intrinsic stress pathways that would be expected to be activated by cytokine exposure, we also exposed INS1 cells to 24 hours of tunicamycin or thapsigargin to model endoplasmic reticulum stress or doxorubicin to induce DNA damage. Tunicamycin (1 µM) (**Figure 4A-C**), thapsigargin (5 nM) (**Figure 4D-F**), and doxorubicin (50 nM) (**Figure 4G-I**) all significantly increased cellular nSMase2, and cellular and EV ceramides.

**Figure 4.**
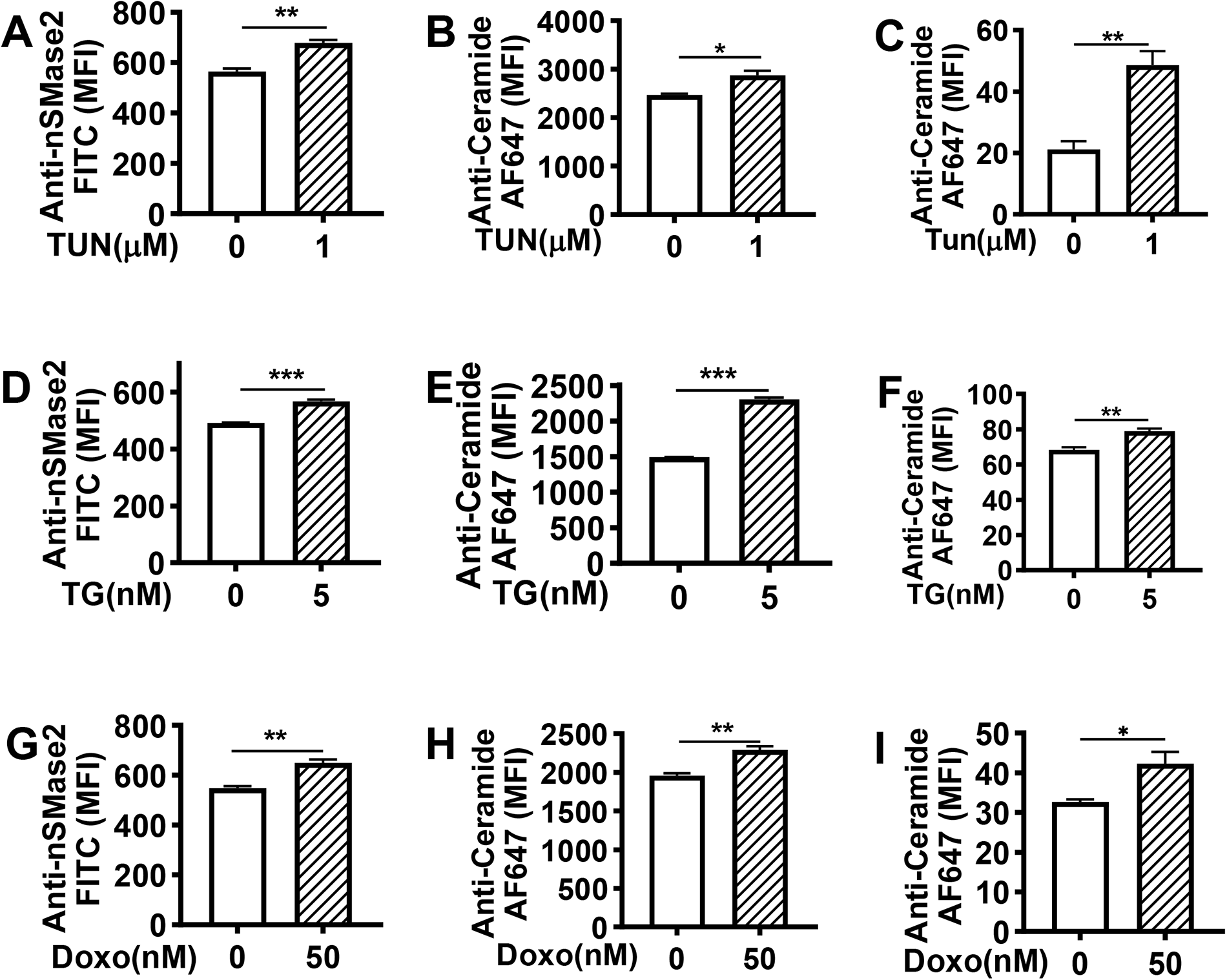
Tunicamycin, thapsigargin, doxorubicin treatment increases β cell nSMase2 and ceramide production in INS1 cells. In INS1 cells, 24 h exposure to (A-C) tunicamycin (1 µM), (D-F) thapsigargin (5 nM), (G-I) doxorubicin (50 nM) increases nSMase2 (A, D, G), ceramides (B, E, H) as well as ceramides in extracellular vesicles (C, F, I). EVs were isolated by immunoaffinity using tetraspanin antibodies. Ceramide content on EVs was determined by flow cytometry using ceramide antibody. N=9, with 3 separate experiments performed. Data expressed as mean ± SEM; *p<0.05; **p<0.01; ***p<0.001

### Ceramide-enriched EVs house distinct miRNA cargo compared to global EV populations

Given our data suggesting changes in ceramide-enriched small β cell EVs after inflammatory cytokine exposure, we next sought to explore the idea that distinct small EV populations enriched for membrane ceramide exist. To test this possibility, we performed immunoaffinity-based small EV pulldowns using antibodies to tetraspanins (CD9, CD63, CD81, global EV membrane proteins) (39) or EV pulldowns using antibodies to ceramide, and performed bulk small RNA sequencing to assay differences in EV cargo, with and without IL1β treatment. To validate the EV pulldown using ceramide antibodies, we captured EVs on magnetic streptavidin microbeads by using ceramide antibody and IgM isotype control, then EVs were stained with Exo-FITC (System Bioscience) and analyzed by flow cytometry (**Supplemental figure 2**).

Here, multidimensional scaling (shown in **Figure 5A**) comparing the small EVs isolated via tetraspanin pulldown vs. EVs isolated via ceramide antibody pulldown suggested that at baseline, significant differences existed in cargo from small EVs isolated via tetraspanin vs. ceramide antibody pulldown, again supporting the idea of ceramide-enriched small EV subpopulations. Changes in miRNA content induced by IL1β treatment were overall similar within both subpopulations. As shown in **Table 1**, ceramide-enriched EVs exhibited significant increases or decreases in expression of at least a 1.5-fold difference of 69 miRNAs (P-value <0.05 and FDR <0.05) compared to global EV populations, with 34 downregulated and 35 upregulated miRNAs. Ingenuity Pathway Analysis identified 3 highly scored networks highly impacted by ceramide-enriched EV miRNAs. The top-scored network included miRNAs and genes involved in insulin signaling (**Figure 5B**). As shown in **Supplemental Figures 3 and 4**, the other highly scored networks included AGO2 (argonaute RISC catalytic component 2), PAX3 (paired box 3), FOXO1 (forkhead box protein O1), and ABCA1 (ATP Binding Cassette Subfamily A Member 1), and AGO2, Argonaute, BRAF(B-Raf proto-oncogene, serine/threonine kinase), CAMTA1 (calmodulin binding transcription activator 1), MTSS1 (MTSS I-BAR domain containing 1), NR0B2(nuclear receptor subfamily 0 group B member 2), NR3C2 (nuclear receptor subfamily 3 group C member 2).

**Figure 5.**
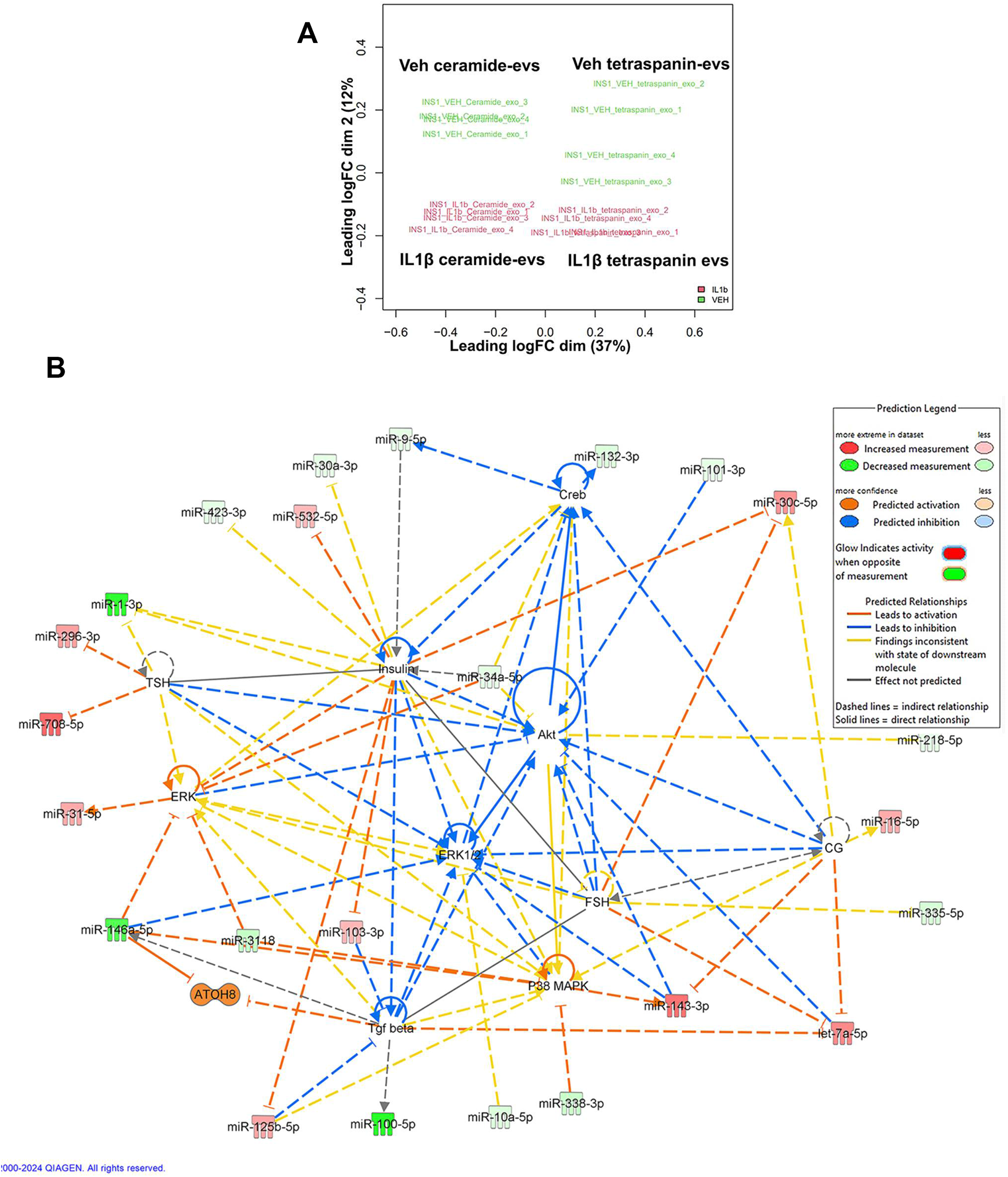
β cell ceramide-enriched EV subpopulations carry differential miRNA cargo compared to global EV populations. INS1 β cells were exposed to IL1β (10 ng/mL) for 24 h. Ceramide enriched EVs and tetraspanin pulldown EVs were isolated using streptavidin magnetic beads and biotinylated antibodies against ceramide or tetraspanins. EV RNA was isolated and subjected to miRNA sequencing. (A) One-dimensional plot of multidimensional scaling (MDS) on miRNA showed baseline differences in miRNA expression profiles between ceramide enriched EVs vs. global EVs. (B) The highest scored network was generated using Qiagen Ingenuity Pathway Analysis (IPA) with a set of cutoffs, down and up 1.5-fold of expression fold change, P-value <0.05 and FDR <0.05. The network suggested that differential miRNAs were involved in a series of network nodes including insulin.

**Table 1.**
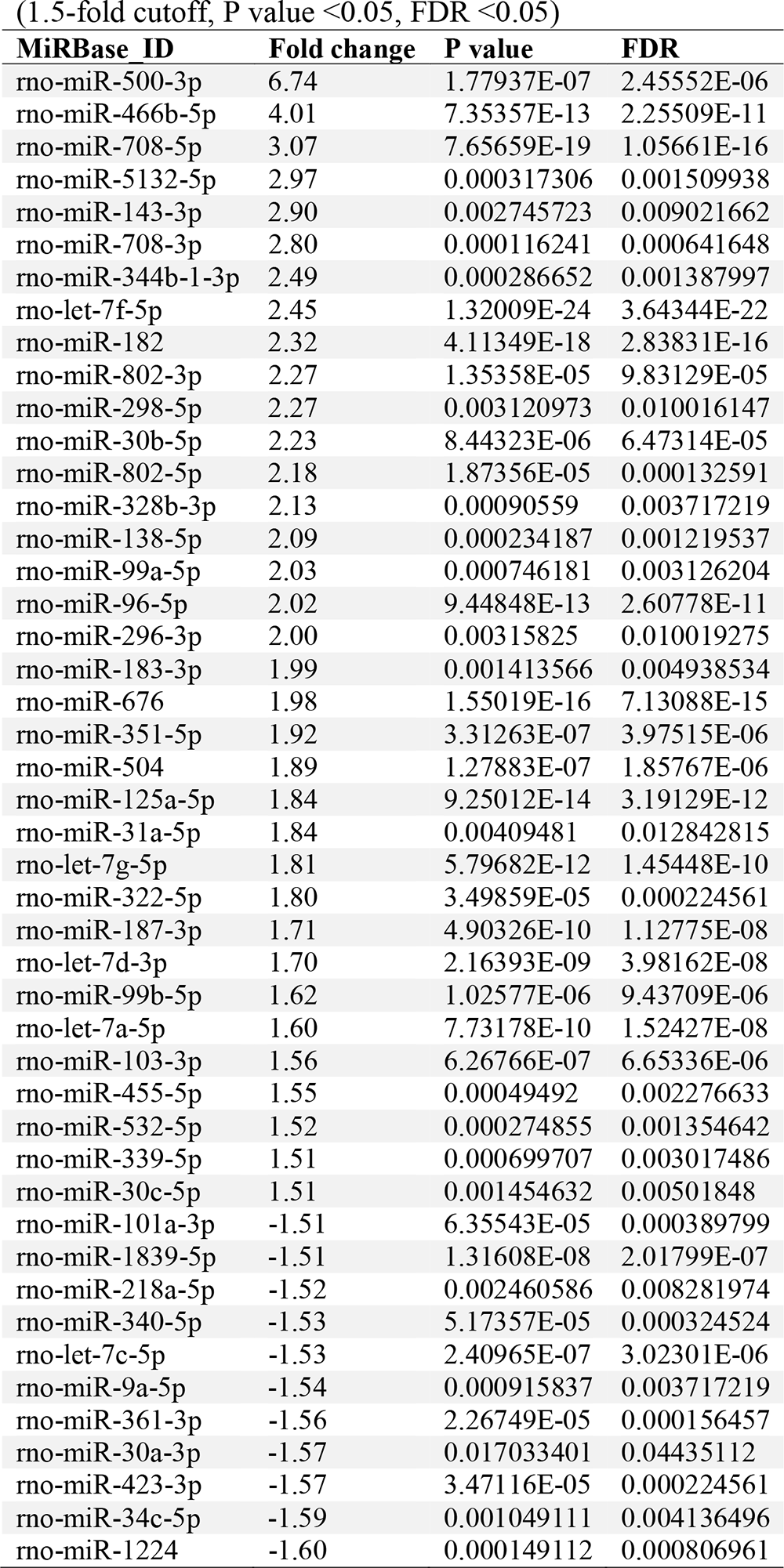
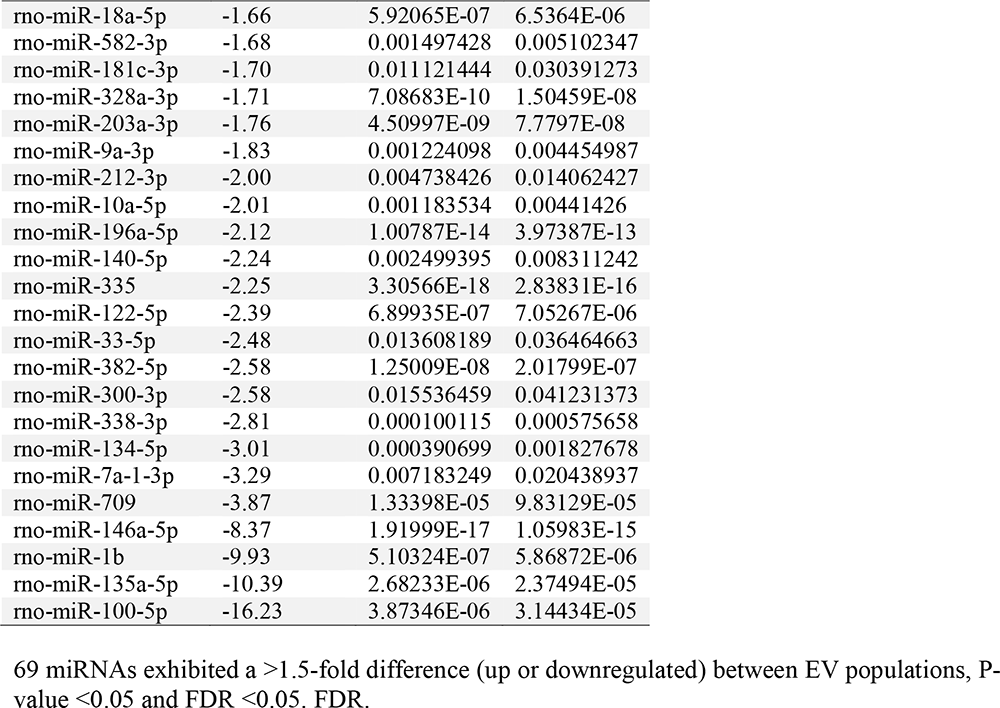
Differential miRNA profiles of ceramide enriched EVs vs. global EVs from INS1 cells.

### β cell EVs can be transferred to other β cells and this process is enhanced by cytokine treatment or chemical activation of nSMase2

Given the potential of EVs to interact with surrounding cells, we asked if activation of the β cell nSMase2/ceramide axis could impact β cell EV transfer. To test this possibility, we utilized INS-1 cells with CD9 (a tetraspanin/EV membrane marker) tagged with GFP (**Supplemental figure 5A**). To quantify EV transfer, CD9-GFP cells were cocultured with INS-1 cells with constitutive mCherry expression for 72 h, then GFP/mCherry double-positive cells were quantified using flow cytometry (**Figure 6A**). At baseline, 1.7% of cocultured cells showed GFP/mCherry double positivity, reflecting CD9-GFP transfer to mCherry cells. Double positivity was significantly increased by IL1β treatment (**Figure 6A and 6C**). A similar increase in CD9-GFP transfer was observed with CAPE treatment to chemically increase nSMase2 activity (**Figure 6B**). To ensure that transfer was not the result of direct cell:cell contact, we also quantified CD9GFP EV transfer into wild-type cells seeded into trans-well inserts (**Supplemental figure 5B**). Here, after either 48 h or 72 h, wild-type INS1 cells within trans-well inserts exhibited GFP fluorescence, suggesting internalization of EVs carrying donor cell GFP.

**Figure 6.**
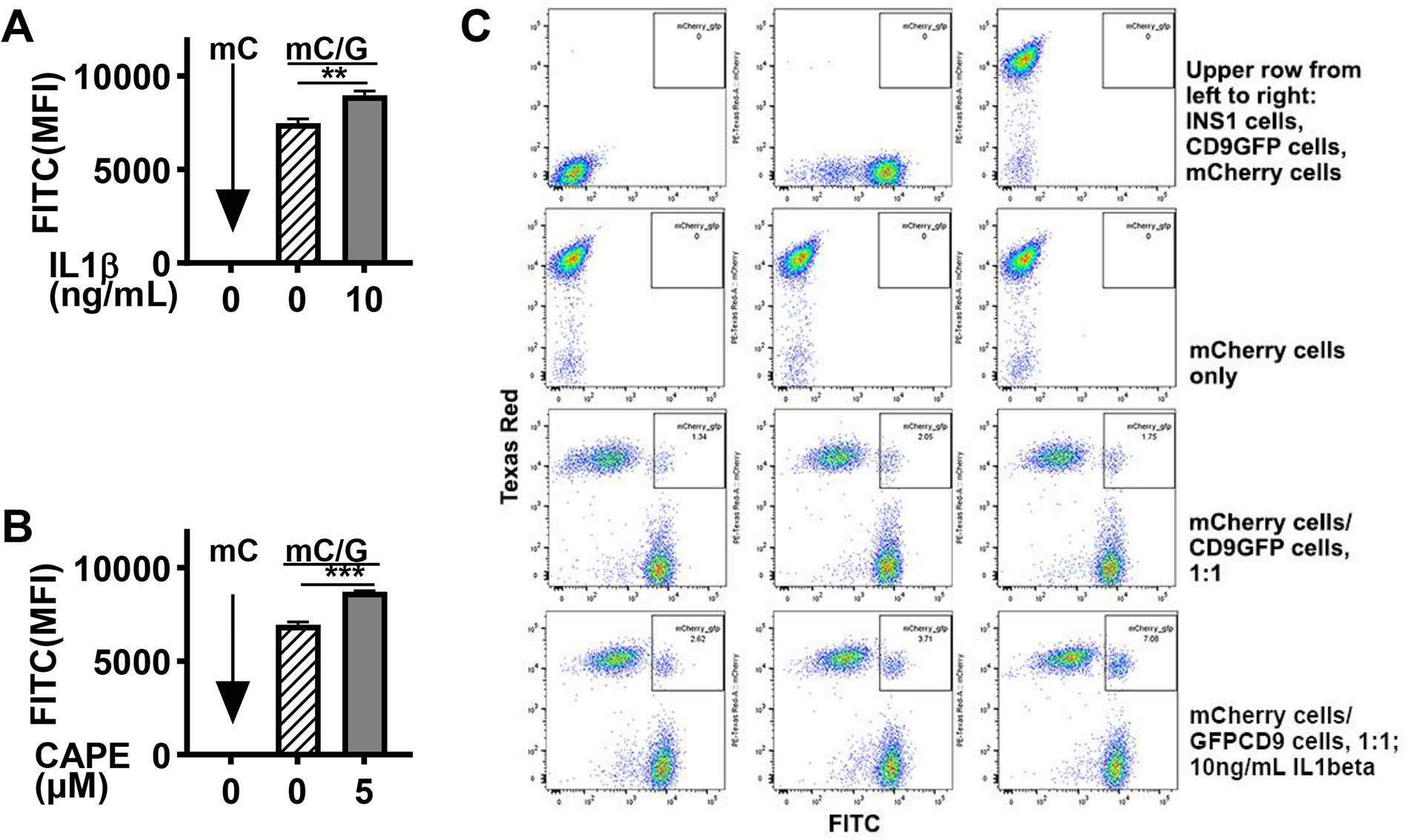
β cell EVs can be transferred to surrounding β cells, and EV transfer is enhanced by IL1 β or CAPE treatment. INS1 β cells harboring GFP tagged CD9 (parent cells) and INS1 cells harboring global mCherry fluorescence (recipient cells) were co-cultured at a 1:1 ratio for 72 h. Quantification of double positive GFP+ mCherry+ cells, reflecting CD9 GFP transfer, was performed using flow cytometry. (A) IL1β treatment (10 ng/mL) significantly increased CD9-GFP EV transfer to neighboring mCherry cells. (B) CAPE (5 µM) treatment also increased EV transfer to neighboring cells. mCherry protein did not transfer to neighbor cells (data not shown). (C) Fluorescent dot plots demonstrate the CD9-GFP EV transfer to neighboring mCherry cells. n=9, with 3 separate experiments performed. .*p<0.05; **p<0.01; ***p<0.001

These findings suggest that not only does cytokine-induced activation of nSMase2 increase ceramide-enriched small EV populations, but that this axis also increases local transfer of β cell EVs to other β cells.

### Total plasma EV ceramide is similar in children with T1D compared to euglycemic pediatric samples

Finally, we asked if ceramide EV content was altered in the circulation of children with type 1 diabetes. We obtained previously bio-banked plasma samples from 26 children with new-onset type 1 diabetes and matched these by age, sex, and BMI to nondiabetic control plasma samples (demographic characteristics in **Supplemental Table 4**). Small EVs were isolated using size exclusion chromatography (**Figure 7A**) and using tetraspanin antibody capture, and EV ceramide was quantified by a ceramide ELISA (**Figure 7B**) and by flow cytometry (**Figure 7C**). We did not detect differences in plasma EV ceramide between new-onset T1D vs. control samples, suggesting that in the context of T1D, cytokine-induced changes in EV ceramide may not be systemic, but rather an organ-specific phenomenon.

**Figure 7.**
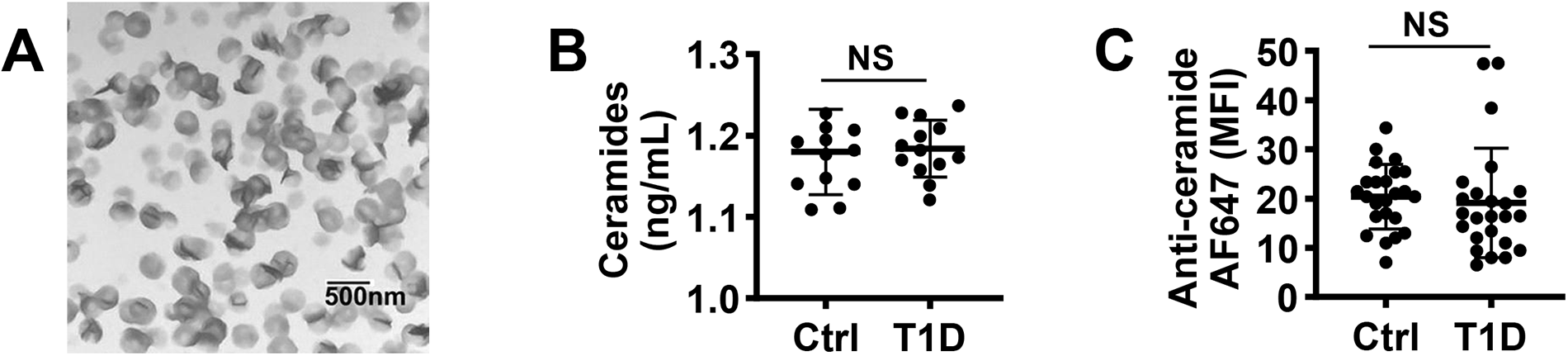
No differences in plasma EV ceramide staining were detected in samples from children with new onset type 1 diabetes (T1D) compared to age matched controls. (A) Human plasma EVs were also isolated by SEC, then lysed using ELISA lysis buffer and diluted 80 times for ceramide ELISA (AGF Bioscience), which also showed no between group differences. (B) Plasma EVs were isolated using tetraspanin bead-based pulldown and ceramides were quantified using flow cytometry. (C) Plasma EVs were isolated by SEC, concentrated and visualized by electronic microscopy. A: 26 donors, B: 48 donors. n=4, with 2 separate experiments performed. Data expressed as mean ± SEM.

## Discussion

Findings from our lab and others have identified changes in β cell EV cargo under disease conditions, and the potential of islet EVs to serve as paracrine effectors, but mechanistic etiologies underlying β cell EV generation and content are poorly understood (6, 7, 9–12, 40, 41). Here we show for the first time, that β cell inflammatory stress yields increases in β cell small EV populations enriched for the bioactive lipid ceramide housing distinct miRNA cargo. Cytokine-induced increases in β cell EV ceramide content were mediated by changes in the expression and activity of nSMase2. Furthermore, inflammatory stress or nSMase2 activation increased β cell EV transfer to surrounding β cells.

Several studies have suggested that islet EVs have potential as paracrine effectors in the islet microenvironment. Cytokine treatment of β cells to induce inflammatory stress induces physiologic changes in EV RNA and protein cargo (9–11, 16, 41). β cells or islet EVs from islets display can display immunostimulatory properties for antigen-presenting cells and CD8+ T cells that are enhanced by cytokine treatment of parent cells (9, 16, 42). Coculture of cytokine treated MIN6 cells led to an miRNA dependent increase in recipient cell apoptosis (10), while CXCL10 on the surface of cytokine treated islet EVs reduced GSIS and expression of identity genes in recipient β cells, while inducing upregulation of chemokine signaling and proteins associated with antigen presentation (16). T lymphocyte EVs can transfer miRNAs to β cells that induce apoptosis and chemokine signaling (9, 43). Islet EVs also stimulate endothelial cell angiogenesis (40) and have the potential to interact with other beta cells.

Small RNA sequencing identified a distinct set of miRNAs in ceramide-enriched EVs that were linked to key pathways involved in beta cell function and identity, including insulin signaling and FoxO1 (44) and miRNA action(45). Our observation that nSMase2 activation is associated with increased β cell EV transfer suggests that certain EV populations or physiologic conditions may potentiate the paracrine effects of transfer or interaction of β cell EVs with surrounding β cells. Future studies will assay the impacts of ceramide-enriched β cell EV populations on other cell types in the islet microenvironment.

In other systems, nSMase2-mediated ceramide synthesis promotes MVE membrane ceramide accumulation, leading to inward vesicle budding and ultimately, release of miRNA-dense, ceramide-enriched EVs (17, 19). Specifically, nSMase2 can be required for the release and transfer of certain exosomal miRNAs (17, 18). Additionally, bioactive EV lipids themselves may potentially be transferred to recipient cells (46). Other mechanisms, such as the ESCRT (endosomal sorting complex required for transport)-dependent pathway, also regulate EV formation and likely contribute to β cell EV formation (47). However, given prior descriptions of ceramide in β cell stress pathways (22–28), a role for changes in in regulation of stress-response-induced regulation of β cell EV content is logical, and our findings suggest that this is in part related to increased nSmase2 expression. Future work will interrogate the role of other regulators of cellular ceramide content and localization (48), and other EV biogenesis pathways.

Testing in plasma did not identify systemic differences in ceramide-enriched EVs in children with new-onset T1D, a condition linked to increased systemic inflammation (49). These findings could suggest organ-specific changes in EV ceramide content in the context of inflammation, rather than a consistent response of all cell types. The improvement in EV ceramide in response to exendin-4 is intriguing, consistent with studies showing that the GLP-1 receptor agonist liraglutide decreased total plasma ceramides in individuals with type 2 diabetes (50) and may represent an additional therapeutic effect of this drug class.

This work had some limitations. EV isolations are well recognized to exhibit heterogeneity (35). We tried to address this by isolating by size and specific membrane components to more selectively isolate subpopulations, and validation of isolations with multiple techniques. However, our preparations likely still contained a combination of EVs derived from the MVE or exosomes and microvesicles, which are classically derived from membrane blebbing (5). Plasma sample analysis was limited by available blood volume, and included systemic EVs from multiple cell types, therefore the relative contribution of β cell EVs was likely limited, especially considering reduced β cell mass in children with T1D.

In aggregate, this work studies provides substantial novel mechanistic data informing understanding of nsMase2-based generation of increased numbers of a subpopulation of ceramide-enriched β cell EVs under condition of inflammatory stress. Compared to global EVs, ceramide-enriched EVs contained distinct miRNA cargo, and increased nSmase2 activity led to increases in β cell EV transfer, suggesting a role in amplification of the paracrine effects of β cell EVs. Targeting and modulation of this process may ultimately allow for novel therapeutics aimed at improving β cell health in diabetes.

## Acknowledgements

This work was partially presented in oral abstract form at the ADA 2022 scientific sessions.

## Author Contributions

JX designed and performed experiments and wrote and edited the manuscript. AH and JE helped perform experiments and edited the manuscript. RGM interpreted experiments and edited the manuscript. EKS designed experiments and wrote and edited the manuscript. All authors agree to the final version of the manuscript. EKS serves as guarantor of this work. This work utilised core services provided by the Diabetes Research Center grant P30 DK097512 (to Indiana University School of Medicine). This work also utilized the Indiana University Flow Cytometry Core and Indiana University Center for Medical Genomics for RNA sequencing analysis, to Nicholas Conoan at University of Nebraska Medical Center Electron Microscopy Core Facility for TEM. The authors thank the members of the Indiana University Melvin and Bren Simon Comprehensive Cancer Center Flow Cytometry Core. The Indiana University Melvin and Bren Simon Comprehensive Cancer Center Flow Cytometry Core is funded in part by NIH, National Cancer Institute (NCI) grant P30 CA082709. Sequencing analysis was carried out in the Center for Medical Genomics at Indiana University School of Medicine, which is partially supported by the Indiana University Grand Challenges Precision Health Initiative. Human pancreatic islets and/or other resources were provided by the NIDDK-funded Integrated Islet Distribution Program (IIDP) (RRID:SCR_014387) at City of Hope, NIH Grant # 2UC4DK098085. **Human islets for research were also provided by the Alberta Diabetes Institute IsletCore at the University of Alberta in Edmonton (**http://www.bcell.org/adi-isletcore.html**) with the assistance of the Human Organ Procurement and Exchange (HOPE) program, Trillium Gift of Life Network (TGLN), and other Canadian organ procurement organizations. Islet isolation was approved by the Human Research Ethics Board at the University of Alberta (Pro00013094). All donors’ families gave informed consent for the use of pancreatic tissue in research.**

