## Supplemental for "Proinflammatory stress activates neutral sphingomyelinase 2 based generation of a ceramide-enriched β cell EV subpopulation"

**Supplemental Figures and Tables**

**Supplemental Figures**

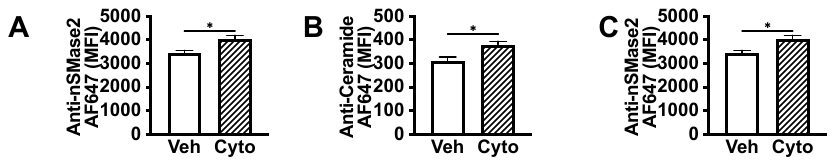

**Supplemental Figure 1. Cytokine treatment increases β cell nSMase2 and ceramide content in MIN6 cells.** In MIN6 β cells, 24 h mix of IL1β (5 ng/mL), TNFα (10 ng/mL) and IFNγ (100 ng/mL) increases (A) nSMase2, (B) total cell ceramides, and (C) cell surface ceramides. N=9, with 3 separate experiments performed.

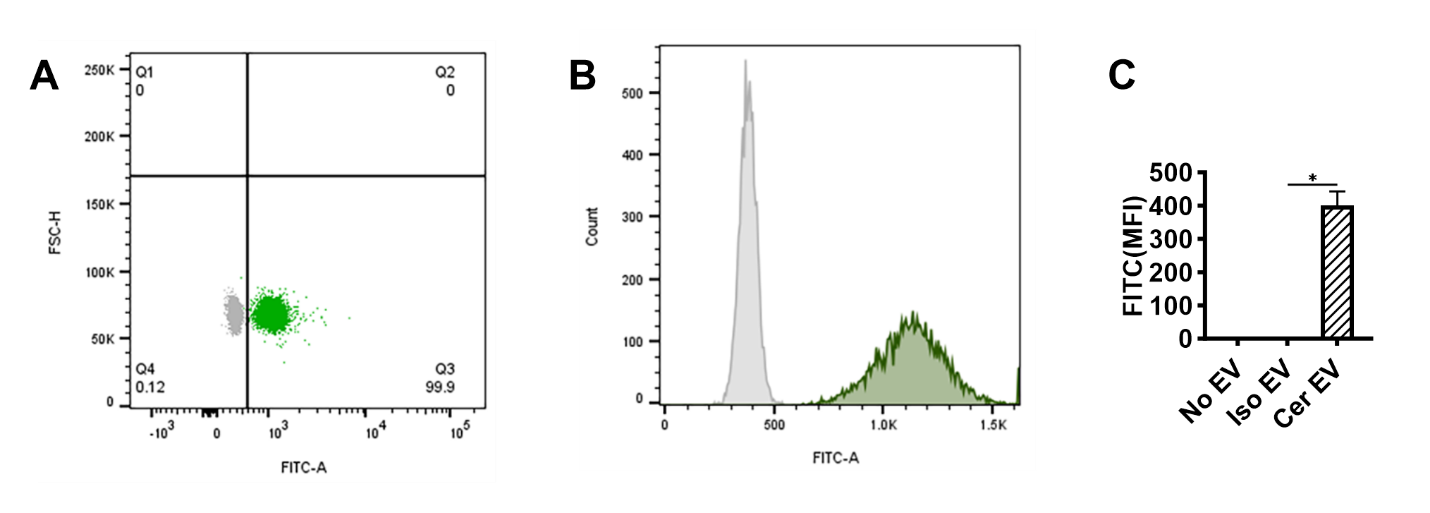
**Supplemental figure 2. Magnetic capture of ceramide enriched EVs.** Ceramide specific EVs were isolated by using coupling streptavidin magnetic beads with biotinylated ceramide antibody following the manual of Exo-Flow exosome capture kit (System Biosciences) that is used to isolate tetraspanin antibody specific EVs. EVs on magnetic beads were stained with Exo-FITC exosome stain (System Biosciences) and analyzed by flow cytometry and FlowJo. (A) Flow cytometry dot plot overlay, (B) histogram plot overlay to compare ceramide specific isolated EVs (Green) compared to isotype control (Grey), and (C) mean fluorescent intensity showing presence of EVs captured by anti-ceramide antibody. No ev=no EV input, but with biotinylated anti-ceramide antibody. Iso= biotinylated isotype control; Cer= EVs isolated using biotinylated anti-ceramide antibody.

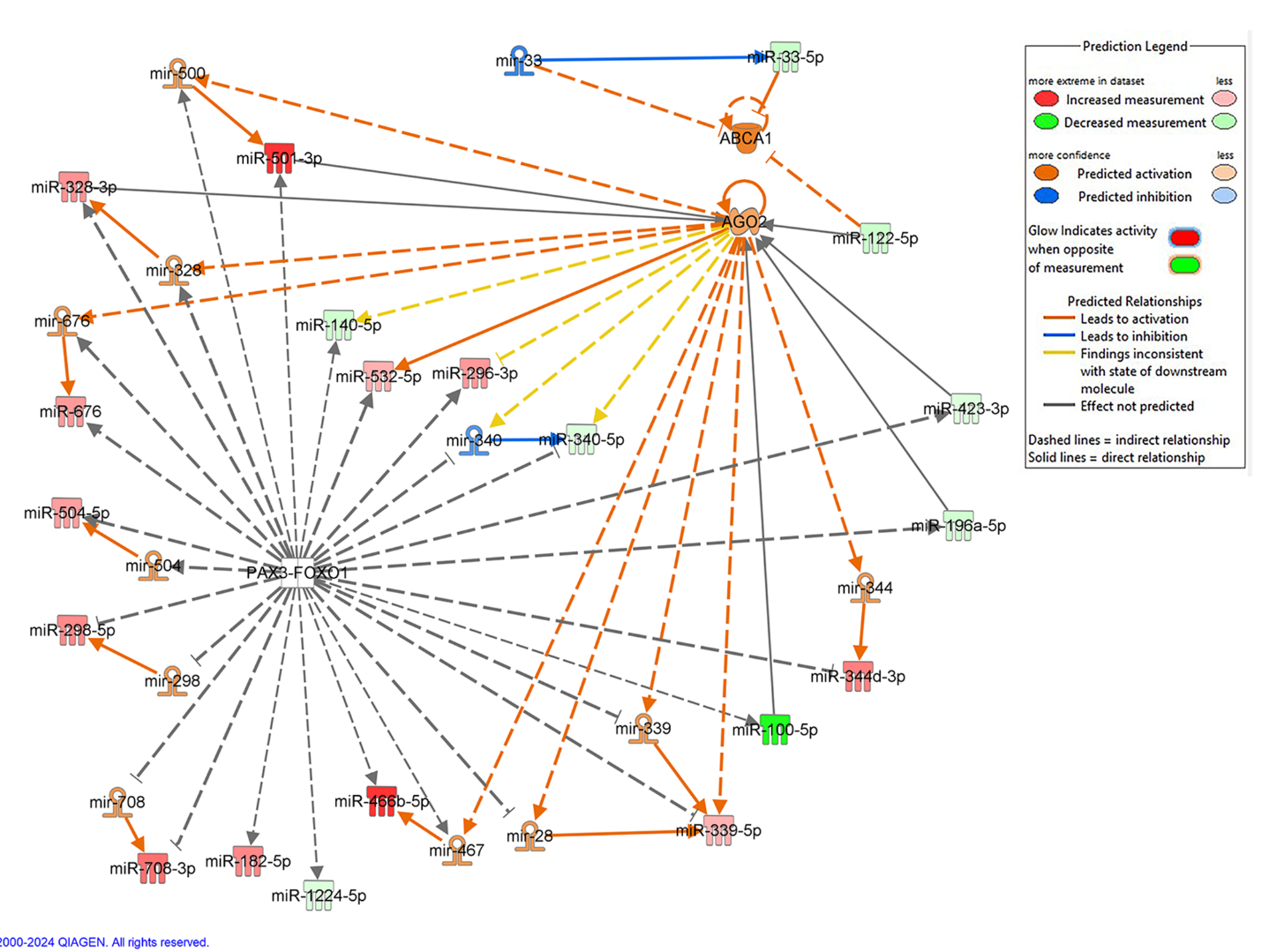

**Supplemental figure 3. Differential miRNA content within ceramide-enriched EVs regulate important signaling nodes.** Ingenuity Pathway Analysis of differential miRNA content within ceramide-enriched versus global EV populations identified 3 highly scored signaling networks, including a network containing AGO2 (argonaute RISC catalytic component 2), PAX3 (paired box 3), FOXO1 (forkhead box protein O1), and ABCA1 (ATP Binding Cassette Subfamily A Member 1) and 32 altered miRNAs.

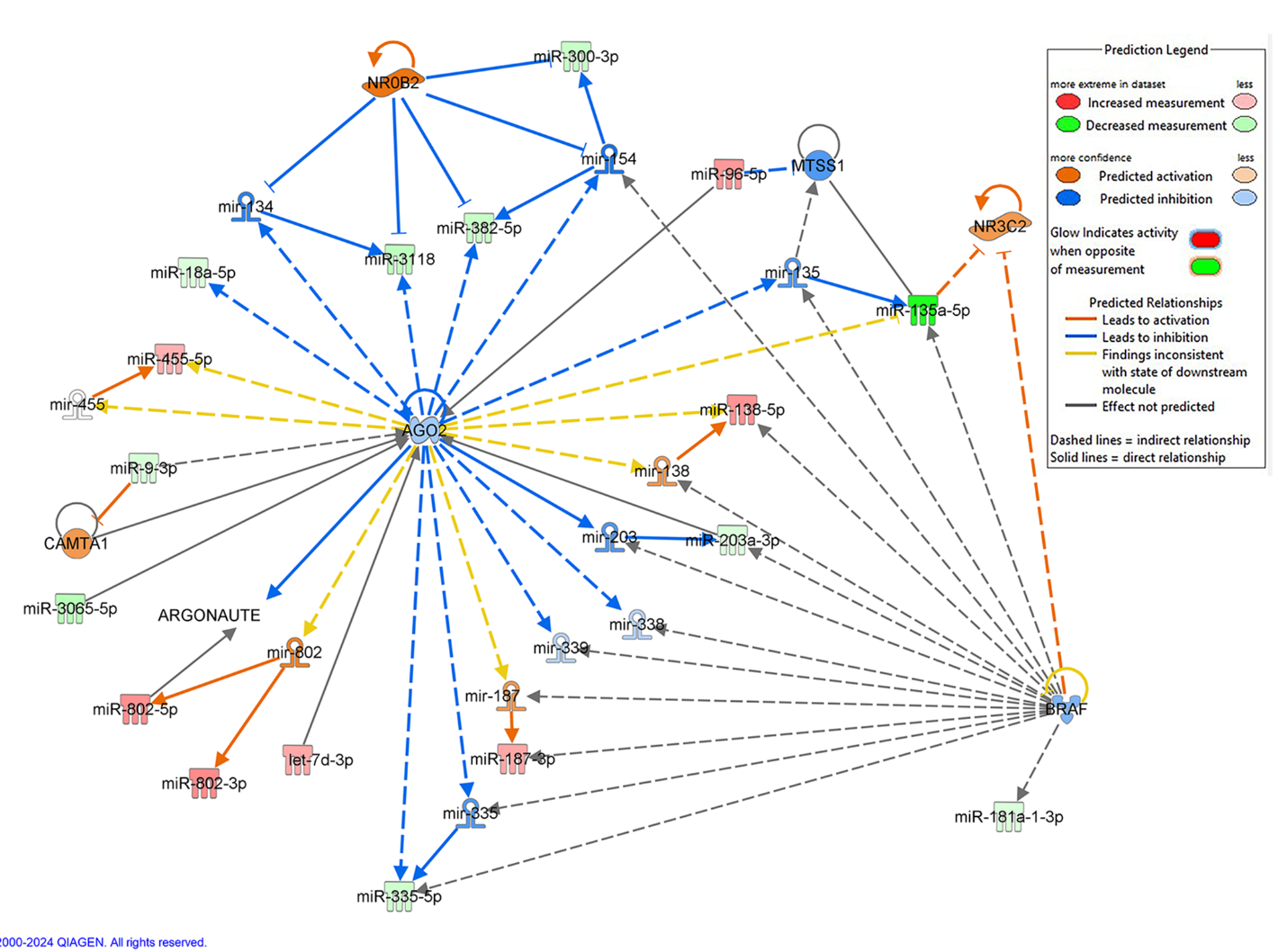

**Supplemental figure 4. Differential miRNA content within ceramide-enriched EVs regulate important signaling nodes.** Ingenuity Pathway Analysis of differential miRNA content within ceramide-enriched versus global EV populations identified 3 highly scored signaling networks, including a network containing AGO2 (argonaute RISC catalytic component 2), Argonaute, BRAF(B-Raf proto-oncogene, serine/threonine kinase), CAMTA1(calmodulin binding transcription activator 1), MTSS1(MTSS I-BAR domain containing 1), NR0B2(nuclear receptor subfamily 0 group B member 2), NR3C2(nuclear receptor subfamily 3 group C member 2), and 28 altered miRNAs.

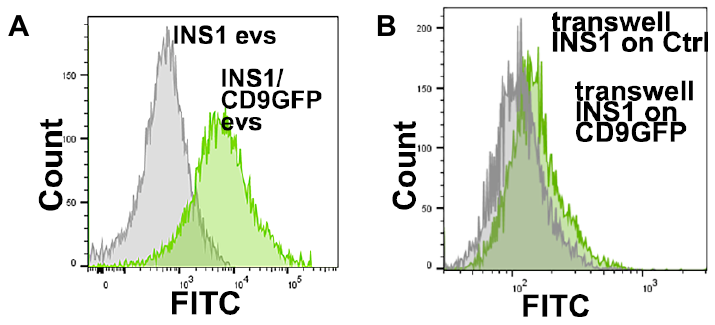

**Supplemental figure 5. EV transfers to neighboring cells.** (A) CD9GFP EVs can be distinguished by flow cytometry from EVs of control INS1 cells. EVs were separated by ultracentrifugation. Grey, INS1 EVs; Green, CD9GFP EVs. (B) CD9GFP EV transfer to recipient cells on trans-wells. Grey, control INS1 cells without CD9GFP EVs; Green, INS1 cells harboring CD9GFP EVs.

**Supplemental Tables**

**Supplemental table 1: Reagent Information**

| Reagents | Vendors |
| --- | --- |
| Exosome depleted FBS | ThermoFisher Scientific |
| IL1β, TNFα, IFNγ | R&D Systems, Minneapolis, MN |
| Thapsigargin | MB Biochemicals, Santa Ana, CA |
| Tunicamycin | MilliporeSigma, Burlington, MA |
| Doxorubicin | MilliporeSigma, Burlington, MA |
| Caffeic acid phenethyl ester (CAPE) | Santa Cruz Bio, CA |
| Exendin-4 | Tocris Bioscience, Bristol, United Kingdom |
| pLJM1-EGFP lentiviral vector, plasmid #19319 | Addgene, Watertown, MA |
| mCherry in pReceiver-Lv213 vector | Genecopoeia, Rockville, MD |
| Rat nSMase2 shRNA with GFP reporter | Origene, Rockville, MD |
| Scrambled shRNA in a pGFP-C-shLenti Vector | Origene, Rockville, MD |
| Biotinylated tetraspanin antibodies (CD9, CD63, CD81) | Novus Biochemicals, Centennial, CO |
| Anti-ceramide antibody clone MID 15B4 | MilliporeSigma, Burlington, MA |
| Exo-Flow Capture kits | System Biosciences, Palo Alto, CA |
| Exosome streptavidin magnetic beads | Invitrogen, Thermo Fisher Scientific |
| Streptavidin magnetic beads | System Biosciences, Palo Alto, CA |
| Exo-FITC exosome stain | System Biosciences, Palo Alto, CA |
| Exosome elution buffer | System Bioscience, Palo Alto, CA |
| qEVsingle /35 nm Legacy column | Izon Science US, Medford, MA |
| Anti-nSMase2 antibody | Santa Cruz Biotechnology, Dallas, TX |
| Anti-β-actin | Cell Signaling Technology, Danvers, MA |
| Goat anti-rabbit IgG (H+L) IRDye 800CW | Li-Cor Biosciences, Lincoln, NE |
| Goat anti-mouse IgG IRDye 680RD | Li-Cor Biosciences, Lincoln, NE |
| Enzyme-free cell dissociation buffer | MilliporeSigma, Burlington, MA |
| 5x ELISA lysis buffer | Cell Signaling Technology, Danvers, MA |
| Ceramide (CER) ELISA Kit | AGF Bioscience, Northbrook, IL |
| miRNeasy Isolation Kits | Qiagen,Germantown, MD |
| RNase-Free DNase Set | Qiagen, Germantown, MD |

**Supplemental table 2. Human islets for islet ceramide and nSMase2 analysis.**

| Donor ID | Sex | Age | BMI | HbA1c | Blood glucose(mg/dl) |
| --- | --- | --- | --- | --- | --- |
| SAMN17928660 | M | 62 | 25.9 |  | 136.4 |
| SAMN18614363 | F | 42 | 30.9 | 5.9 | 140.2 |
| SAMN18718846 | M | 46 | 33.5 | 5 | 104.6 |
| SAMN20923891 | M | 45 | 50.6 | 5.4 | 150.2 |
| SAMN25860453 | M | 44 | 25.8 | 6 | 133.2 |
| SAMN29208584 | M | 38 | 29.5 | 5.2 | 107 |

Human islets were provided from the Integrated Islet Distribution Program (IIDP) at City of Hope and The Alberta Diabetes Institute IsletCore.

**Supplemental table 3. Human islets for islet EV ceramide analysis.**

| Donor ID | Sex | Age | BMI | HbA1c | Blood glucose(mg/dl) |
| --- | --- | --- | --- | --- | --- |
| SAMN30648329 | M | 54 | 21.5 | 5.2 | 105.8 |
| SAMN30927410 | M | 50 | 36.3 | 4.9 | 240.8 |
| SAMN31242271 | M | 52 | 37.5 | 5.3 | 181 |
| SAMN31645456 | M | 68 | 30.9 | 5.4 | 195.2 |
| SAMN32641506 | M | 16 | 29.5 | 5.4 | 216.4 |
| SAMN33902448 | F | 32 | 19.1 | 4.6 | 112.6 |
| R483 | M | 36 |  |  |  |

Human islets were provided from the Integrated Islet Distribution Program (IIDP) at City of Hope and The Alberta Diabetes Institute IsletCore.

**Supplemental table 4. Donor demographics for plasma analyses**

| Group | Control | T1D |
| --- | --- | --- |
| Age | 9.2 | 8.9 |
| Female | 38.5% | 38.5% |
| BMI | 18.3±0.9 | 17.2±0.8 |

Plasma samples were obtained from 26 children with T1D and 26 age, sex, and BMI-matched controls ranging from plasma at the age of 3 to 15. No significant differences in age, BMI, or sex distribution were present between groups.
